# β-Lactam antibiotics trigger mouse TLR2/4- and human TLR2/5-dependent Jarisch-Herxheimer reaction in leptospirosis

**DOI:** 10.64898/2026.04.29.721589

**Authors:** Stylianos Papadopoulos, Thomas Bernard, Dorian Joffres, Frédérique Vernel-Pauillac, Julie Cagliero, COSIPOP study group, Catherine Werts

**Affiliations:** Institut Pasteur, Université Paris Cité, INSERM U1306, Unité de Biologie et Génétique de la Paroi Bactérienne, Paris, F-75015, France; École Normale Supérieure de Lyon, Department of Biology, Lyon, France; École Normale Supérieure Paris-Saclay, Department of Biology, Université Paris-Saclay, Gif-sur-Yvette, France; Institut Pasteur of New Caledonia, Institut Pasteur International Network, Leptospirosis Research and Expertise Unit, Nouméa, New Caledonia; Institut Pasteur, Centre d’investigation clinique “Investigation clinique et volontaires en santé humaine” (INVOLvE), Paris, -F-75015, France

**Keywords:** One health, Leptospirosis, Spirochetes, β-Lactam antibiotics, Jarisch-Herxheimer reaction, Inflammation, Cytokine storm, Toll-like receptors, Innate immunity, Human blood, Cytokines, Amoxicillin, Ceftriaxone

## Abstract

Leptospirosis is a neglected zoonotic disease causing ∼1 million cases and 60,000 deaths annually. Antibiotic therapy can trigger a detrimental inflammatory Jarisch-Herxheimer reaction (JHR). We investigated antibiotic effects in leptospirosis, using a mouse model of severe disease and human whole blood from healthy donors infected ex vivo with bioluminescent *Leptospira interrogans*. We tested the effect of amoxicillin and ceftriaxone, two bactericidal β-Lactams antibiotics and assessed bacterial survival, cytokines, pathophysiology and JHR mechanisms upon amoxicillin. β-lactams induced profound pro-inflammatory cytokine release. In mice, the inflammation was TLR2 and TLR4 dependent, accordingly to the TLR4 host-specificity of LPS recognition, and amoxicillin exacerbated disease severity within hours, notably worsening myocarditis. In human whole blood, this inflammation depended largely on TLR2 (lipoprotein receptor) and TLR5 (flagellin receptor). Interestingly, only stealthy virulent isolates of *Leptospira interrogans,* unlike some *Leptospira* species belonging to intermediate and non-virulent clades such as *biflexa*, induced in human whole blood a JHR-like effect, which was also observed with *Borrelia burgdorferi*. These findings demonstrate that host-specific TLR recognition of released spirochaetal components by antibiotic treatment drives JHR, evoking a cytokine storm and worsening outcomes.

## Introduction

Leptospirosis is a neglected globally distributed zoonotic infection caused by pathogenic spirochetes of the genus *Leptospira*. Despite causing an estimated one million cases and nearly 60 000 deaths annually, leptospirosis is underdiagnosed and underreported, particularly in low- and middle-income countries where access to diagnostics and surveillance is limited^1^. Climate change, flooding, and rapid urbanisation are expected to further increase disease incidence by enhancing environmental persistence of *Leptospira* and human exposure^2^. Clinical manifestations range from mild febrile illness to severe disease with multiorgan failure, including Weil’s disease and pulmonary haemorrhagic syndrome^3^. Leptospirosis is commonly considered as a disease leading to inflammation, since pro-inflammatory cytokines are measured in patients^4^. However, experimental and recent clinical studies consistently show low levels of circulating pro-inflammatory cytokines at admission, even in severe disease^5,6^. This suggests that leptospirosis may not be a hyperinflammatory condition which is consistent with *in vitro* studies. Indeed, a defining feature of pathogenic *Leptospira* is their remarkably stealthy interaction with the host immune system. This immune silence is consistent with leptospiral multiple immune evasion strategies, including poor recognition by Toll-like receptors (TLRs) and Nod-like receptors^7^. Human TLR4 fails to recognise leptospiral lipopolysaccharide (LPS) because of its atypical lipid A structure, leaving TLR2, the receptor of lipoproteins the primary innate sensor of *Leptospira*^8,9^. In parallel, leptospires actively inhibit inflammasome-driven pyroptosis, an inflammatory cell death leading to massive IL-1β release, further limiting inflammation amplification^10^. These mechanisms minimising early inflammation may favour systemic dissemination and disease progression.

However, this apparent paradox of *Leptospira* being stealthy despite being known to trigger inflammation is highlighted by a recent clinical study of leptospirosis in New Caledonia which demonstrated that the initiation of antibiotic therapy triggered a sudden inflammatory burst in approximately 50% of patients, associated with a Jarisch-Herxheimer reaction (JHR)^6^.

JHR is an acute inflammatory syndrome occurring within hours of antibiotic initiation and characterised by fever, chills, rigors, hypotension, and transient clinical deterioration^11^. JHR is now recognised as a general feature of spirochaetal infections, including syphilis, Lyme borreliosis, and relapsing fever, rather than a pathogen-specific phenomenon^12,13^. Despite more than a century of clinical observation since Jarisch and Herxheimer first described acute inflammatory worsening after antibiotic treatment of syphilis^14,15^, it remains uncertainties about JHR in terms of pathogenesis, clinical significance, and management. Early clinicians regarded JHR as a “therapeutic hazard” inherent to spirochaetal infections, attributed to rapid bacteriolysis and endotoxin release^11,16^. However, as our immunological understanding has become more detailed, the biological understanding of JHR has shifted towards models that emphasise the activation of the innate immune system by specific bacterial components that have not yet been identified.

In leptospirosis, like in other spirochaetoses, current treatment guidelines from WHO recommend the use of β-lactams especially in severe disease, but other antibiotics such as macrolides or tetracyclines are also mentioned^17^. Of note β-lactam antibiotics, including penicillin, amoxicillin, and ceftriaxone, exert bactericidal activity by inhibiting cell wall synthesis^18^, leading to rapid bacterial disruption of the bacteria, and the release of pro-inflammatory bacterial components. By contrast, macrolides, which inhibit protein synthesis^19^ and are bacteriostatic, could potentially prevent such rapid cell wall disruption.

In this study, we investigated the mechanisms underlying the JHR in leptospirosis treated with ß-lactam antibiotics using the complementary *in vitro* human whole blood model and an *in vivo* mouse model of lethal infection^20^.

## Results

### JHR-like inflammatory effect is induced by amoxicillin

Previously, we demonstrated in a clinical study that patients with leptospirosis exhibit low or undetectable levels of pro-inflammatory cytokines, including IL-1β, IL-6, and TNF, upon admission to the hospital, although they exhibit high levels of the anti-inflammatory IL-10 cytokine^6^. Conversely, after antibiotic treatment, 50% of leptospirosis patients exhibited clinical signs of JHR and was associated with increased levels of IL-6, TNF and IL-10^6^. To further study this phenomenon *in vivo*, we used a mouse model of acute, lethal leptospirosis, which we previously described^5,20^. We injected the mice at the peak of leptospiral dissemination on day 3 post-infection (pi) with 100 μg/mouse of amoxicillin, the same weight-dose administered to patients in New Caledonia. Three hours after the antibiotic injection, at the peak of inflammation observed in human patients^6^, mice were euthanised and blood collected for cytokine dosage (Fig. 1A).

**Figure 1:**
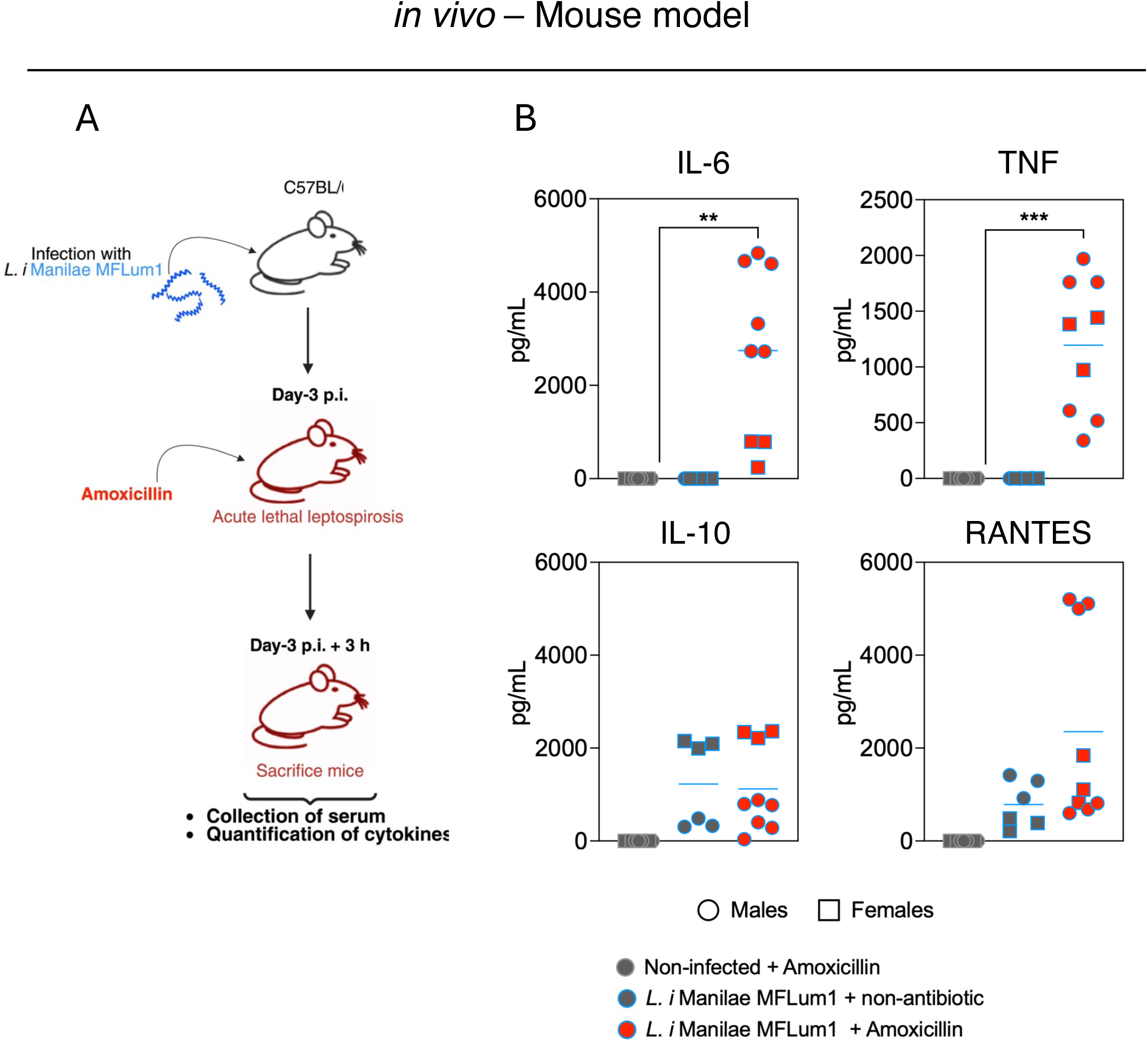
**Amoxicillin-induced JHR effect *in vivo in mice*** A) Schematic representation of the *in vivo* protocol used to assess JHR following amoxicillin administration in mice infected with *Leptospira interrogans* serovar Manilae strain MFLum1. *B)* Serum cytokine concentrations (IL-6, TNF, IL-10, and RANTES). Grey symbols with grey outline represent uninfected control mice treated with amoxicillin plus hydrocortisone; grey symbols with blue outline represent infected untreated mice; red symbols with blue outline represent infected mice treated with full-dose amoxicillin; half-red/half-blue symbols represent infected mice treated with gradual amoxicillin; half-red/half-dark blue symbols represent infected mice treated with gradual amoxicillin plus hydrocortisone. Data are mean of n=6-9 mice pooled from three independent experiments. Statistical analyses were performed using Welch’s t test. Open circles represent male mice, and open squares represent female mice. Significance threshold was *p* ff 0.05 (\**p* ff 0.05, \*\**p* ff 0.01, \*\*\**p* ff 0.001).

As previously shown, in absence of amoxicillin, IL-6 and TNF were undetectable when amoxicillin was not injected (Fig. 1B, grey symbols with a light blue border), however, after its administration, amoxicillin led to striking increased levels of IL-6 and TNF (Fig. 1B, red symbols). As previously observed, the infection led to increased levels of the RANTES chemokine^6^ that were further increased upon antibiotic. Conversely, elevated levels of the anti-inflammatory IL-10 cytokine were observed as previously shown upon *Leptospira* infection, but the antibiotic treatment did not modify these levels. No elevation of cytokines was found in uninfected control mice treated with amoxicillin, suggesting that the antibiotic itself was not responsible for the inflammation.

### JHR-like effect aggravates the pathophysiology of the mice *in vivo*

To further study the physiopathology consequences of antibiotic treatment and JHR-like effect, we measured the serum amyloid A (SAA), an acute-phase protein and inflammatory marker comparable to the C-reactive protein (CRP) in humans. Levels of SAA, that are increased in infected mice as previously shown^5^, were significantly higher in mice who received amoxicillin (Fig. 2A). Similarly, we quantified serum LDH, a marker of lytic cell death (Fig. 2B). Consistent with our previous results^5^, serum LDH levels increased with infection, and levels were significantly higher when mice were injected with amoxicillin (Fig. 2B). Then we tested the vascular permeability since lethal leptospirosis is associated with increased vascular leakage^5^. To this end, we performed an Evans blue assay on MFLum1-infected individuals who were or were not injected with amoxicillin. After perfusion of the mice, the livers, lungs, and kidneys were collected, and the dye was extracted from the tissues to measure the optical density. Our results showed that, upon amoxicillin injection, vascular permeability increased slightly in the livers (Fig. 2C). Similarly, although not significant, there was a tendency for higher levels of Evans blue in the lungs and kidneys of mice that received the antibiotic. As previously reported^5^, lethal leptospirosis is associated with myocarditis. To further characterise the extent of pathophysiological deterioration, we quantified circulating levels of cardiac troponin I (cTnI), a sensitive and specific biomarker of myocardial injury. Consistent with these findings, amoxicillin treatment in severely infected individuals resulted in a twofold increase in cTnI levels (Fig. 2D), indicating exacerbation of cardiac injury. Histopathological analysis further supported these findings, revealing increased infiltration of F4/80⁺ macrophages and CD3⁺ T cells in cardiac tissue following amoxicillin treatment, consistent with aggravated myocarditis (Fig. 2E). No difference was observed in non-infected mice that were or were not injected with amoxicillin. No discrepancy was detected between male and female mice in any of the parameters evaluated. Overall, injecting amoxicillin into lethally infected mice with leptospirosis led to JHR-like effect, which worsens the pathophysiology.

**Figure 2:**
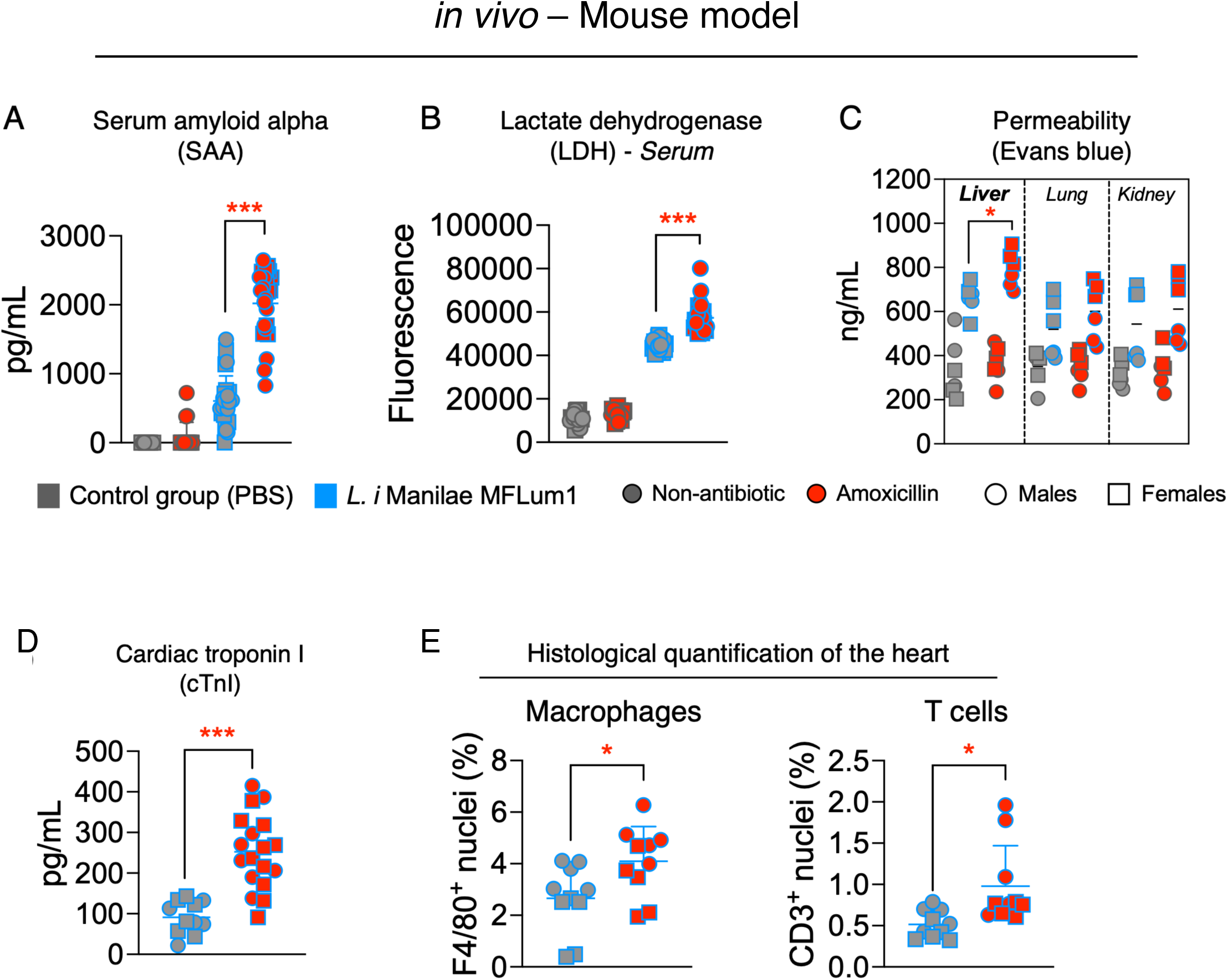
**β-lactam-induced JHR effect exacerbates systemic inflammation and tissue injury *in vivo*** Assessment of systemic inflammation and tissue injury following amoxicillin administration in *Leptospira interrogans* serovar Manilae strain MFLum1-infected mice. (A) Serum amyloid A (SAA) concentrations as a marker of acute-phase inflammation. Data are mean of n=17-24 mice pooled from seven independent experiments. (B) Serum lactate dehydrogenase (LDH) activity as an indicator of tissue injury. Data are mean of n=16-24 mice pooled from seven independent experiments. (C) Vascular permeability assessed by Evans blue dye extravasation in liver, lung, and kidney. Data are mean of n=6 mice pooled from three independent experiments. Grey symbols with grey outline represent uninfected untreated mice; grey symbols with blue outline represent infected untreated mice; red symbols with grey outline represent uninfected mice treated with amoxicillin; red symbols with blue outline represent infected mice treated with amoxicillin. (D) Cardiac troponin I (cTnI) as a marker of myocarditis. Data are mean of n=11 MFLum1-infected mice without antibiotic treatment, and n=18 MFLum1-infected mice treated with amoxicillin pooled from 5 independent experiments. (E) Quantification of F4/80^+^ macrophages and CD3^+^ T cells in the hearts of *L. interrogans* Manilae MFLum1-infected mice treated or not with amoxicillin. Data are mean of n=10 mice pooled from two independent experiments. Blue bars with grey symbols represent infected untreated mice; blue bars with red symbols represent infected mice treated with amoxicillin. For panels A, B, D and E: grey symbols with grey fill represent uninfected untreated mice; grey symbols with red fill represent uninfected mice treated with amoxicillin; blue symbols with grey fill represent infected untreated mice; blue symbols with red symbols fill infected mice treated with amoxicillin. Statistical analyses were performed using Welch’s t test. Open circles represent male mice, and open squares represent female mice. Significance threshold was *p* ff 0.05 (\**p* ff 0.05, \*\**p* ff 0.01, \*\*\**p* ff 0.001).

### Amoxicillin-induced JHR-like effect involves TLR2 and TLR4 in mice

We then investigated in mice the innate immune receptors involved in the phenomenon of JHR effect induced by amoxicillin. Because amoxicillin weakens the bacterial cell envelope, we hypothesised that this would expose LPS, lipoproteins and other immunogenic structures, such as flagellins. Depending on host specificity, these leptospiral structures subsequently signal through TLR2 and TLR5 in human, or TLR2 and TLR4 in mice^7–9,21,22^. To validate these results *in vivo*, we used transgenic mice deficient in TLR2 or TLR4, following the previously described protocol with MFLum1 infection and amoxicillin injection. As previously demonstrated^23^, *Tlr4^-/-^* mice exhibited increased susceptibility to infection, as evidenced by more severe clinical signs and greater weight loss compared to *Tlr2^-/-^* and wild-type (wt) mice (Sup. Fig. 1). Regarding the cytokine response a statistically significant decrease in IL-6, TNF and RANTES was measured in both *Tlr2^-/-^* and *Tlr4^-/-^* mice, suggesting that production of these pro-inflammatory cytokines upon amoxicillin treatment mostly depends both on TLR2 and TLR4 receptors, with the chemokine RANTES mostly dependent on TLR4 (Fig. 3B), as already observed^23^. No sex differences were found in mice. Overall, the amoxicillin-induced cytokine production depended on both TLR2, the lipoprotein receptor and TLR4, the LPS receptor.

**Figure 3:**
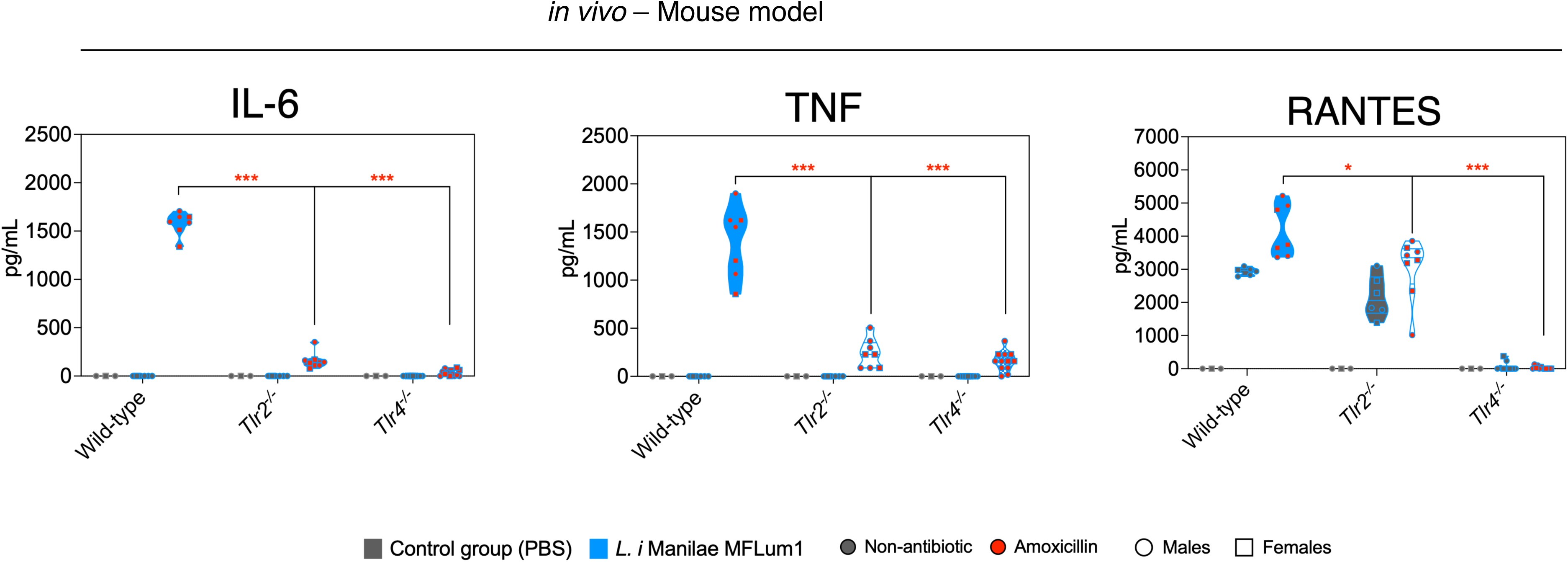
TLR2/4-dependent mechanisms underlying JHR effect in mice. Serum cytokine concentrations (IL-6, TNF, and RANTES) in wild-type, *Tlr2^-/-^*, and *Tlr4^-/-^*mice following amoxicillin administration. Grey symbols indicate infected untreated mice; red symbols indicate infected mice treated with amoxicillin. Coloured violin plots represent wild-type mice; white violin plots with blue outline represent *Tlr2^-/-^* mice; patterned violin plots represent *Tlr4^-/-^* mice. Data are mean of n=7-12 mice pooled from three independent experiments. Statistical analyses were performed using Welch’s t test. Open circles represent male mice, and open squares represent female mice. Significance threshold was *p* ff 0.05 (\**p* ff 0.05, \*\**p* ff 0.01, \*\*\**p* ff 0.001).

It would have been interesting to test whether bacteriostatic antibiotics, such as azithromycin (a macrolide) and doxycycline (a tetracycline), which inhibit protein synthesis without targeting the bacterial cell wall and are used to treat leptospirosis, would also trigger this inflammation. To this end, we conducted a preliminary experiment to compare the time course of antibiotic efficacy *in Leptospira* infection in RAW 264.7 murine macrophages. This involved measuring bioluminescence at various time points after antibiotic administration to assess leptospiral killing. Amoxicillin and ceftriaxone, a third-generation β-lactam also commonly used to treat leptospirosis, killed all the leptospires within three hours of treatment (Sup Fig. 2A). This time frame corresponds to the time at which amoxicillin’s effects were studied in mice^5^ and have been measured in patients with leptospirosis^6^. In contrast, azithromycin and doxycycline both failed to kill leptospires after 3 and 6 hours of treatment, although they were effective at 24 hours (Sup Fig. 2A). Failure to kill *Leptospira* within the three-hour timeframe prevented further testing of their effects on inflammation and comparison with β-lactam antibiotics in our JHR model.

### β-lactams are responsible for the JHR-like effect *in vitro* in human whole blood

To further study the mechanisms of JHR in humans, we investigated whether amoxicillin and ceftriaxone, could give rise to the JHR-like effect *in vitro*. First, whole blood from healthy human volunteers was *in vitro* infected with two doses (10^7^ and 10^6^) of the MFLum1 strain. Then, to mimic an ongoing infection, we administered in each blood donor at 5 hours post-infection (hpi) a dose of antibiotic (Fig. 4A). At 24 hpi, we collected the contents of the wells for cytokine dosage and bacterial survival testing. In most donors, the levels of IL-1β, IL-6, and TNF increased upon amoxicillin treatment (Fig. 4B), likewise *in vivo* in mice. Similar results were obtained upon ceftriaxone treatment (Fig. 4C). As expected, leptospiral viability measured by bioluminescence totally decreased after antibiotic treatment, demonstrating the efficacy of both antibiotics against *Leptospira* in human whole blood (Fig. 4D). These results were consistent across both infection doses with MFLum1. No difference was observed between male and female donors (open circles for males and open squares for females). Taken together, these results demonstrate that the use of β-lactams, such as amoxicillin and ceftriaxone in whole blood infected *in vitro* with *L. interrogans* Manilae led to a high secretion of pro-inflammatory cytokines characteristic of JHR observed in patients, such as IL-6 and TNF. However, although some blood donors untreated by antibiotics exhibit low levels of IL-1ß when infected, for most donors leptospiral infection of whole blood triggers the IL-1ß secretion, unlike that observed in leptospirosis patients’ blood^6^.

**Figure 4:**
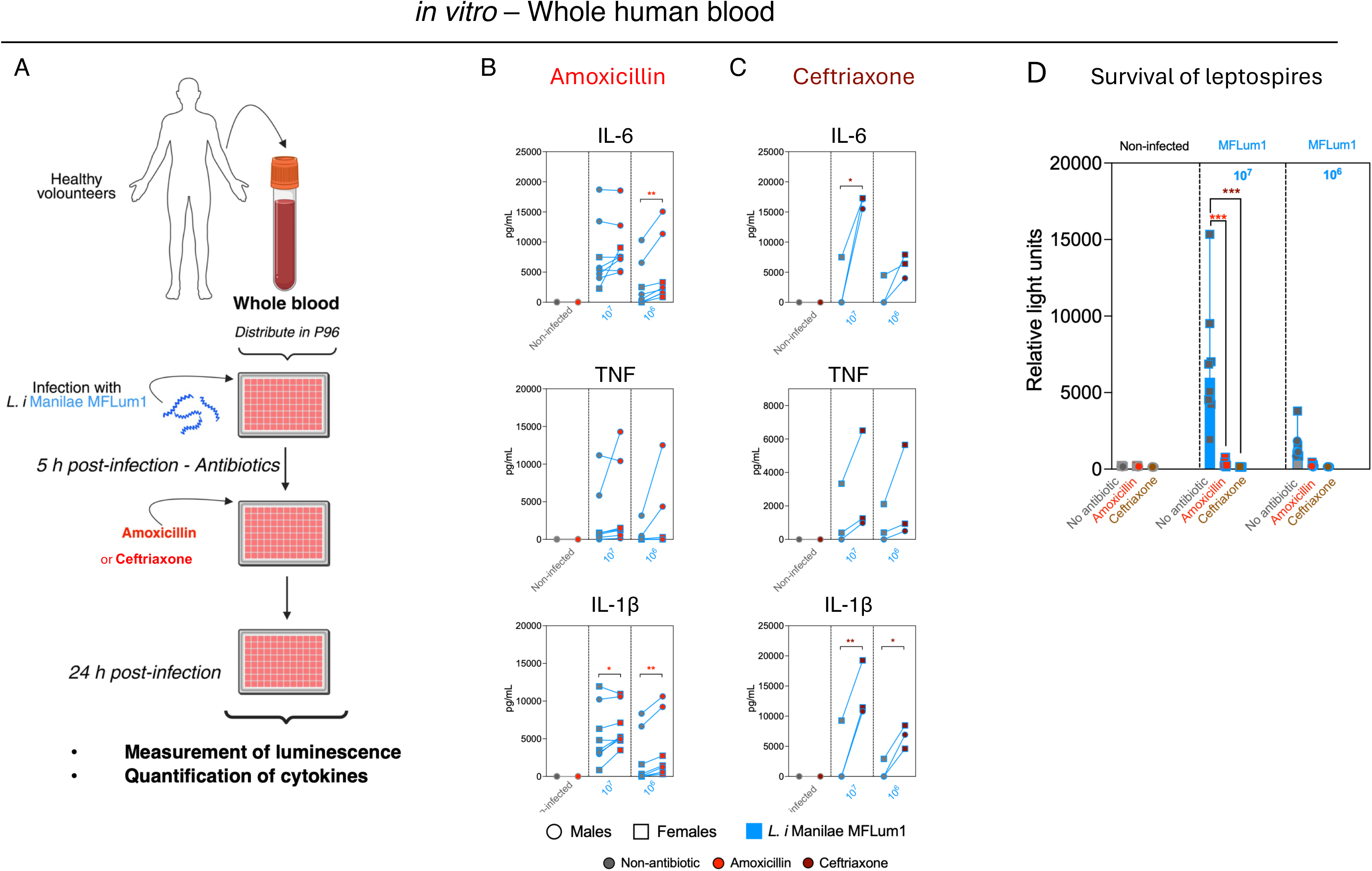
β-lactam antibiotics induce JHR-associated cytokine responses in human blood. (A) Schematic representation of the in vitro protocol used to assess JHR following amoxicillin administration in human whole blood infected with *Leptospira interrogans* serovar Manilae strain MFLum1. (B) Cytokine concentrations (IL-6, TNF, and IL-1β) in uninfected whole blood, or infected with 10^6^ or 10^7^ or bacteria with or without amoxicillin treatment. (C) Cytokine concentrations (IL-6, TNF, and IL-1β) in uninfected whole blood, or infected with 10^6^ or 10^7^ bacteria with or without ceftriaxone treatment. Only significant comparisons between infected untreated and treated groups are shown. Open circles represent male donors, and open squares represent female donors. (D) Survival of leptospires in blood 24h post infection measured by bioluminescence of MFLum1. (B-C) Data are mean of n=6-7 donors (amoxicillin) pooled from six to seven independent experiments, and n=3 donors (ceftriaxone) pooled from three independent experiments. Statistical analyses were performed using two-way ANOVA with Šidák’s multiple comparison test (factor 1: technical replicates per donor; factor 2: antibiotic regimen). Only significant comparisons between infected untreated and treated groups are shown. Open circles represent male donors, and open squares represent female donors.

### Amoxicillin-induced JHR-like effect involves TLR2 and TLR5 in human whole blood

We then investigated in humans the innate immune receptors involved in the phenomenon of JHR effect induced by β-lactams treatments. Therefore, we first incubated human whole blood with neutralising antibodies against TLR2, TLR5 and TLR4, before infection *in vitro* with *L. interrogans* Manilae MFLum1 and further treatment with amoxicillin, as previously described. To control for the efficiency of neutralisation, we used the main agonists of the TLRs: Pam3CSK4 for TLR2, pure LPS of *Escherichia coli* for TLR4, and flagellin of *Pseudomonas aeruginosa* for TLR5. Our results showed an efficient and specific TLR neutralisation with all antibodies in whole blood (Sup Fig. 3). Upon TLR2 neutralisation, after infection with *L. interrogans* and antibiotic treatment, levels of all pro-inflammatory cytokines, including IL-1β, IL-6, and TNF, were significantly lower than in non-neutralised or isotype control conditions (Fig. 5). This suggested that most of the inflammation due to JHR is dependent on TLR2. These results were consistent with previous studies showing that in humans TLR2 is the main receptor involved in leptospires recognition due to the abundance of lipoproteins^8^. Similarly, neutralising TLR5 also led to a decrease in cytokine levels, indicating that this TLR is also involved in the JHR effect, which is also consistent with previous studies showing that in human whole blood, leptospires were recognised by TLR5^24^, and that human TLR5 is able to recognise degraded leptospires, in contrast to murine TLR5^21^. Conversely, neutralisation of TLR4 did not affect cytokine levels (Fig. 5), confirming previous studies showing that human TLR4 does not recognise LPS from *Leptospira interrogans*^8^ in contrast to murine TLR4 that recognises leptospiral LPS but only through the MyD88 pathway^9,22^.

**Figure 5:**
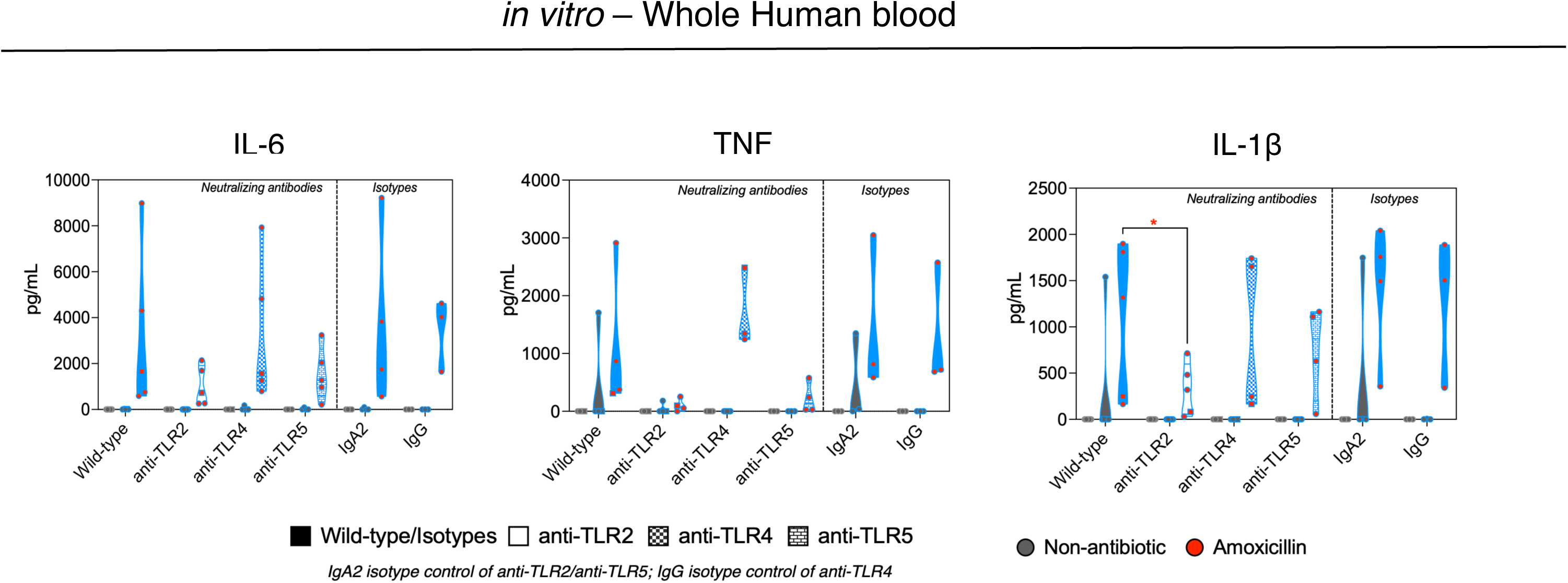
TLR2/5-dependent mechanisms underlying JHR effect in human whole blood. *In vitro* cytokine concentrations (IL-6, TNF, and IL-1β) in human whole blood following amoxicillin treatment in the presence of neutralising antibodies against TLR2, TLR4, or TLR5, or corresponding isotype controls (IgA2 and IgG). Grey symbols indicate infected blood without antibiotic treatment; red symbols indicate amoxicillin-treated blood. Coloured violin plots represent wild-type or isotype control conditions; white violin plots with blue outline indicate TLR2 neutralisation; patterned violin plots indicate TLR4 or TLR5 neutralisation as specified in the panel. Data are mean of n=3-5 donors pooled from three to five independent experiments. Statistical analyses were performed using two-way ANOVA with Šidák’s multiple comparison test (factor 1: technical replicates per donor; factor 2: neutralising antibody or isotype condition). Only significant comparisons between infected untreated and treated groups are shown. Open circles represent male donors, and open squares represent female donors.

No sex differences were found *in vitro* using human blood. Overall, the amoxicillin-induced cytokine production associated with the JHR effect in humans and mice largely depended on TLR2, the lipoprotein receptor.

### β-lactam induced JHR-like effect depends on leptospiral virulence and is species-specific

Next, we investigated whether the JHR-like effect was restricted to *L. interrogans* serovar Manilae or extended to other serovars, and species. First, we infected human whole blood *in vitro* with two serovars initially isolated from human patients: *L. interrogans* Icterohaemorrhagiae (Verdun Vir Cl3) and Copenhageni (Fiocruz LV2756) (Fig. 6A)(Table 1). We then assessed the impact of β-lactam antibiotics (amoxicillin and ceftriaxone). Both antibiotics efficiently cleared leptospires, as confirmed by growth assay (Sup. Fig. 2B). We next quantified pro-inflammatory cytokines. Despite expected inter-donor variability, both serovars consistently triggered robust increases in IL-1β, IL-6 and TNF following β-lactam treatment (Fig. 6A).

**Figure 6:**
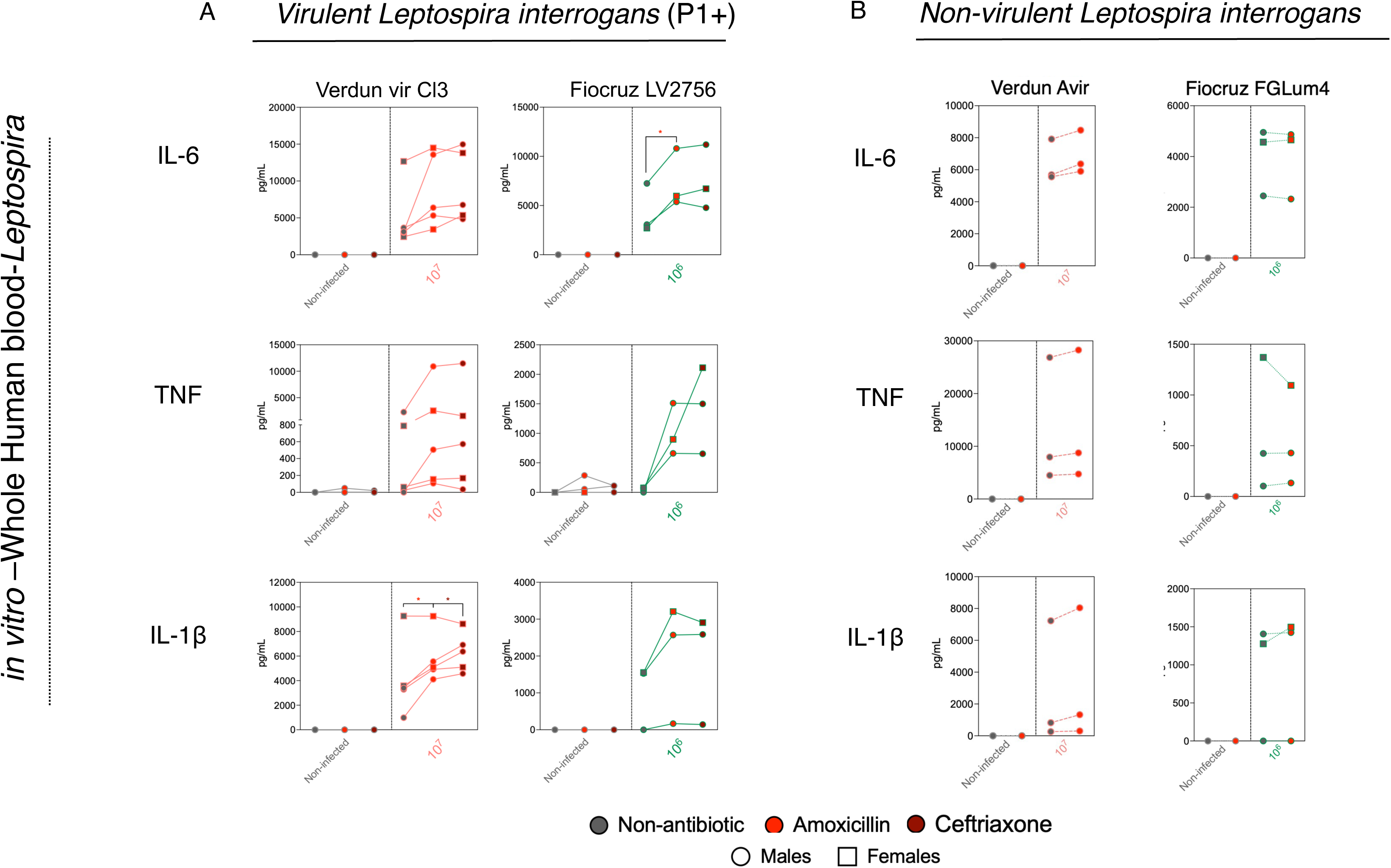
Species-dependent variation in JHR-associated inflammatory responses. Comparison of cytokine concentrations (IL-6, TNF, and IL-1β) in whole blood infected infected with different *Leptospira* serovars and species following antibiotic administration. (A) Virulent P1+ *Leptospira interrogans* serovar Copenhageni strain Fiocruz LV2756 (red) and Icterohaemorrhagiae strain Verdun Cl3 (green) and treated with amoxicillin (red symbols), or ceftriaxone (dark red symbols). (B) Non-virulent strains of highly passaged Icterohaemorrhagiae Verdun Avir (green) and Copenhageni strain Fiocruz FGLum4 (red) and treated with amoxicillin (red symbols), Data are mean of n=2-5 donors (Fiocruz), n=3 donors (Verdun & Patoc) and pooled from two to five independent experiments as indicated in each panel. Statistical analyses were performed using two-way ANOVA with Šidák’s multiple comparison test (factor 1: technical replicates per donor; factor 2: antibiotic regimen). Only significant comparisons between infected untreated and treated groups are shown. Open circles represent male donors, and open squares represent female donors. Significance threshold was *p* ff 0.05 (\**p* ff 0.05, \*\**p* ff 0.01, \*\*\**p* ff 0.001). Each line corresponds to a different donor.

**Table 1.** ***Leptospira* and *Borrelia* species used for infections.**

| Species (subclade) | Serovar | Strain | Origin | Virulence (Animal) |
| --- | --- | --- | --- | --- |
| *Leptospira interrogans*<br>(P1+) | Manilae | MFLum1 | Human, L495 bioluminescent derivative | Yes (kills mice) |
|  | Icterohaemorrhagiae | Verdun Vir Cl3 | Human | Yes (kills guinea pig), No (mice) |
|  |  | Verdun Avir<br>(Cl3, p104) | Human, Lab derivative of Verdun Vir Cl3 | No (guinea pig, mice) |
|  | Copenhageni | Fiocruz LV2756 | Human | Yes (mice) |
|  |  | Fiocruz FGLum4 | Human, L1-130 bioluminescent derivative | No (mice) |
| *Leptospira adleri*<br>(P1-) |  | FH2-B-D1 | Environment<sup>1</sup> |  |
| *Leptospira licerasiae*<br>(P2) | Varillal | VAR 010 | Rat<sup>2</sup> | Yes (rat) |
| *Leptospira biflexa*<br>(S1) | Patoc | Patoc I<br>PFLum7 | Environment<br>Environment, bioluminescent derivative | No |
| *Leptospira kobayashii*<br>(S2) |  | E30 | Environment<sup>3</sup> | No |
| *Borrelia burgdorferi* | B31-A3 |  |  | Yes |

- 1 Thibeaux, R. *et al.* Biodiversity of Environmental *Leptospira*: Improving Identification and Revisiting the Diagnosis. *Frontiers in Microbiology* **Volume 9 - 2018** (2018). https://doi.org:10.3389/fmicb.2018.00816 - 2 Matthias, M. A. *et al.* Human leptospirosis caused by a new, antigenically unique *Leptospira* associated with a *Rattus* species reservoir in the Peruvian Amazon. *PLoS Negl Trop Dis* **2**, e213 (2008). https://doi.org:10.1371/journal.pntd.0000213 - 3 Nakao, R., Masuzawa, T., Nakamura, S. & Koizumi, N. Complete Genome Sequence of *Leptospira kobayashii* Strain E30, Isolated from Soil in Japan. *Microbiology Resource Announcements* **10**, e00907-00921 (2021).

Next, as these strains are pathogenic *L. interrogans*, we sought to assess the role of virulence in the JHR-like effect. We therefore repeated the experiments using a non-virulent derivative of the Icterohaemorrhagiae serovar (Verdun Avir), obtained after 104 *in vitro* passages, and FGLum4 a non-virulent bioluminescent derivative of strain Copenhageni Fiocruz (Table 1). These strains did not induce an increase in cytokine production following β-lactam treatment (Fig. 6B).

Finally, because *L. interrogans* belongs to highly pathogenic P1+ clade among more than 60 *Leptospira* species, we investigated whether this phenomenon extended to other clades. We therefore tested *L. adleri* (P1-) and *L. licerasiae* (P2), which are less pathogenic and typically associated with mild disease, as well as the non-pathogenic saprophytic species *L. biffexa* Patoc (S1) and *L. kobayashii* (S2) (Table 1). Notably, none of these species induced a JHR-like effect following amoxicillin treatment (Sup. Fig. 4) Overall, our results, indicate that β-lactam-induced JHR-like effect was species-dependent. Among pathogenic *L. interrogans* P1+ serovars, only low-passage, highly virulent strains induced a robust JHR effect in human blood, whereas non-virulent derivatives, intermediate species, and saprophytic leptospires did not.

### β-lactam-induced JHR-like effect due to *Borrelia burgdorferi*

We expanded our study to test whether JHR effect also occurred in the same manner with other spirochetes, all characterised by their high content in lipoproteins and endoflagella. We repeated the experimental procedure described previously, infecting human whole blood *in vitro* with two doses (10^7^ and 10^6^) of the spirochete *Borrelia burgdorferi* B31-A3, which causes borreliosis, or Lyme disease. We administered the antibiotics amoxicillin and ceftriaxone at 5 hpi and then, at 24 hpi we quantified the pro-inflammatory cytokines IL-1β, IL-6, and TNF, as previously done with *Leptospira*. *Borrelia* induced the JHR effect upon β-lactam administration, which led to significantly increased levels of all three pro-inflammatory cytokines (Fig. 7). No obvious differences were observed between male and female blood donors.

**Figure 7:**
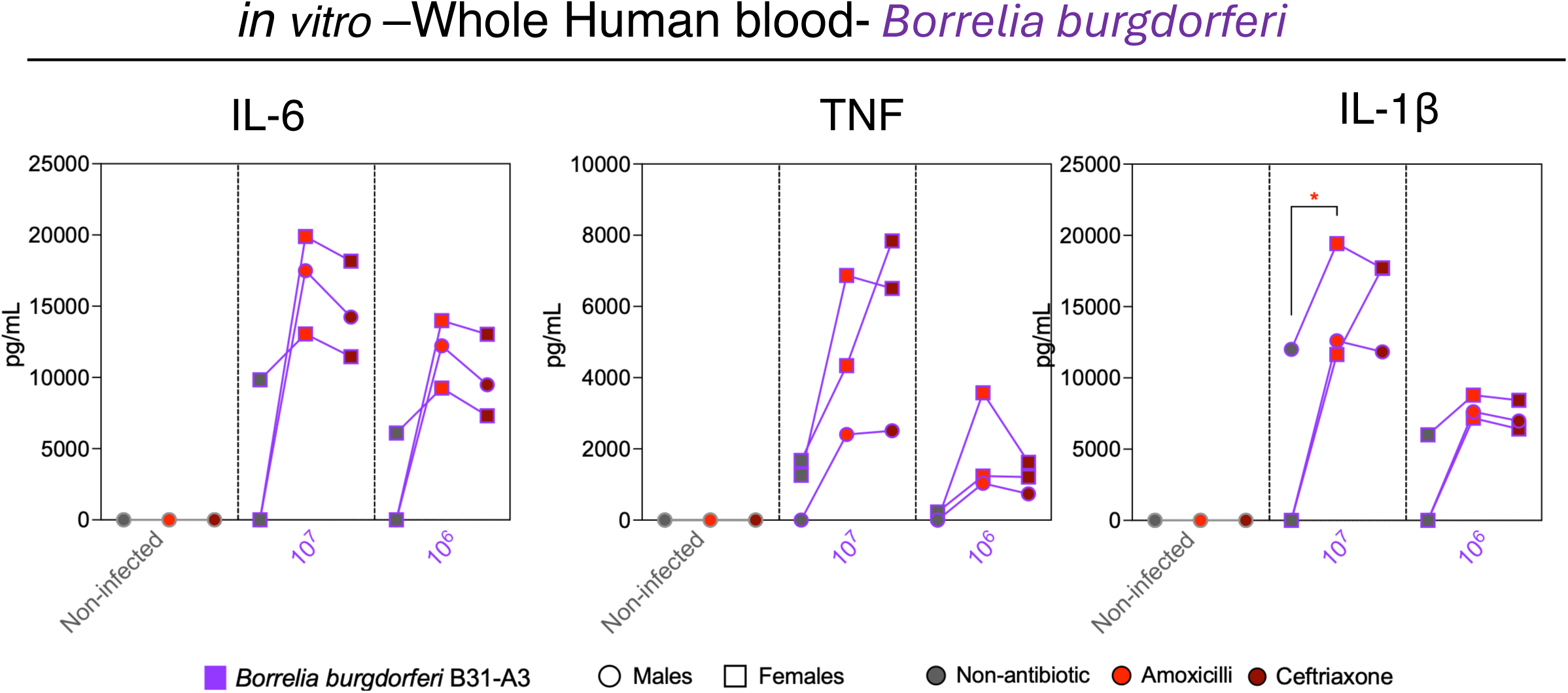
β-lactam-associated cytokine responses in *Borrelia burgdorferi* infection. *In vitro* cytokine concentrations (IL-6, TNF and IL-1β) in human whole blood infected with 10^7^ or 10^6^ *Borrelia burgdorferi* sensu stricto strain B31-A3 following antibiotic administration. Red symbols indicate amoxicillin; dark red symbols indicate ceftriaxone. Data are mean of n=3 donors pooled from three independent experiments. Statistical analyses were performed using two-way ANOVA with Šidák’s multiple comparison test (factor 1: technical replicates per donor; factor 2: antibiotic regimen).

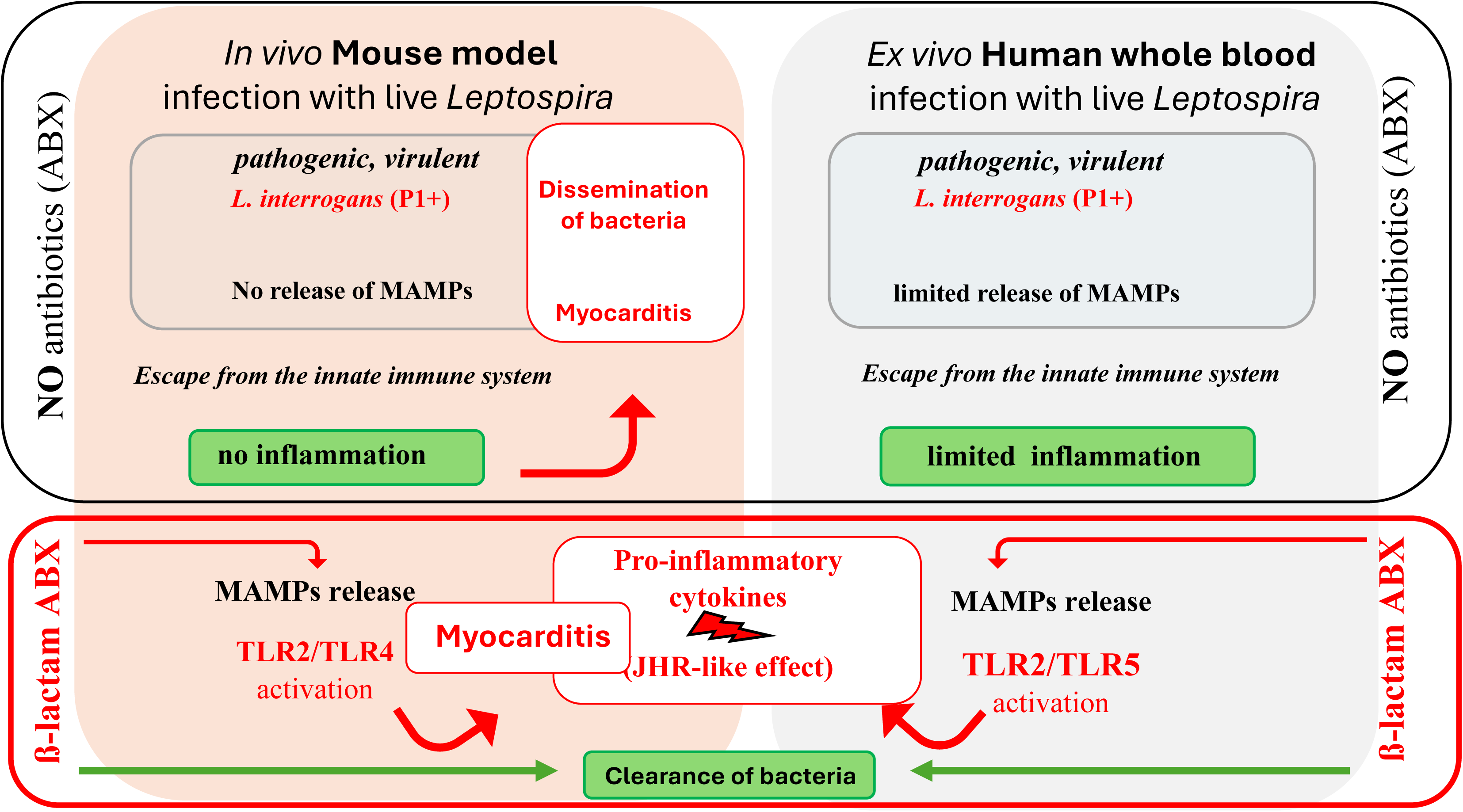

## Discussion

Taken together, our results obtained from *Leptospira* and *Borrelia* reinforce a unified model in which β-lactam antibiotics induce rapid structural disruption of spirochetes, leading to abrupt exposure of lipoproteins and other immunostimulatory components. These molecules engage pattern-recognition receptors (PRRs), particularly human TLR2 and TLR5, and trigger a cytokine-driven inflammatory cascade, characterised by rapid induction of TNF, IL-6, IL-1β, and chemokines. This response may vary according to leptospiral host-related virulence. Importantly, we demonstrated a clinical significance of JHR, as we show aggravated leptospirosis in β-lactam treated-mice. In addition, our experimental findings provide a mechanistic framework for the clinical observations recently reported^6^, who described the JHR in leptospirosis as a clinically relevant, treatment-associated inflammatory deterioration. These findings provide mechanistic support for future studies exploring targeted immunomodulation to prevent JHR-associated morbidity in leptospirosis.

In the context of leptospirosis, JHR has historically been underreported and often dismissed as a mild, transient febrile response. However, clinical cohorts in endemic regions, such as New Caledonia, a French overseas territory and in the southwest Pacific Ocean, have documented significant systemic inflammation, hypotension, and organ dysfunction temporally associated with antibiotic treatments, mostly β-lactam antibiotics, in 10 to 50% of patients with leptospirosis^6,25,26^. Our study aimed to clarify the JHR phenomenon in leptospirosis. We demonstrated *in vitro* in human blood and *in vivo* in mice that β-lactam antibiotics, such as amoxicillin and ceftriaxone, specifically trigger an increased innate immune response, characterised by the secretion of pro-inflammatory cytokines (IL-1β, IL-6, TNF) cytokines, known to amplify leukocyte recruitment and endothelial activation. While species-specific differences in innate immune recognition exist, the overall concordance between *in vitro* human blood data and *in vivo* mouse findings strengthens the translational relevance of this model.

Importantly, our *in vivo* experiments demonstrate that β-lactam-induced JHR-like effect is not merely an inflammatory phenomenon but is harmful to the host. The mouse model of acute lethal leptospirosis used in this study closely recapitulates key features of severe human leptospirosis, including systemic dissemination, (non-)inflammatory responses, vascular leakage, and organ involvement, including the heart^5^. Using this mouse model, we showed that amoxicillin administration at the peak of leptospiral dissemination rapidly led to increased vascular permeability, and elevated levels of acute-phase and tissue-damage markers, including SAA and LDH. In addition, we also provided evidence of exacerbated cellular infiltrates characteristic of myocarditis, accompanied by increased circulating levels of cTnI, suggesting that β-lactam-induced bacterial disruption rapidly aggravates cardiac injury, with measurable effects occurring within hours. These findings are particularly significant given that leptospirosis has been repeatedly associated with widespread endothelial dysfunction, capillary leakage, and haemorrhagic manifestations, as documented in studies of severe pulmonary involvement and vascular injury in human leptospirosis^27,28^ as well as often overlooked myocarditis^29^. Although extrapolation from animal models should be made with caution, our data raise the possibility that similar early and treatment-associated exacerbation of cardiac injury may occur in patients with leptospirosis receiving β-lactam antibiotics. By exacerbating TNF- and IL-6-dependent endothelial injury, JHR effect may actively worsen organ damage rather than accompany severe disease. These pathological changes can occur without overt differences in gross clinical parameters^12,13^, highlighting that the true extent of JHR-associated injury may be underestimated when relying solely on traditional clinical markers. The present study demonstrates that β-lactam antibiotics can worsen leptospirosis. This finding could help clarify the inconsistent benefits of antibiotics observed in human leptospirosis systematic studies^30^. Specifically, antibiotic treatment appears to be effective when administered prior to the onset of organ failure, but not in patients with established organ dysfunction^31,32^. Indeed, the inflammatory adverse JHR may potentially mask the beneficial effect of β-lactams in clearing leptospires.

Although IL-1β is frequently measured as a marker of inflammation in spirochaetal infections, accumulating evidence indicates that IL-1β plays a limited functional role in leptospirosis. *L. interrogans* LPS actively blocks pyroptosis in human and murine cells, thereby uncoupling IL-1β production from downstream inflammatory cell death pathways^10^. In this context, our findings support a model in which leptospiral JHR pathology is driven predominantly by TNF- and IL-6-dependent signalling rather than by IL-1β-mediated inflammasome activation. However, it is important to reconcile the discrepancy between IL-1β dynamics in clinical leptospirosis and in controlled cellular infection assays. In clinical *in vivo* studies of leptospirosis patients exhibiting JHR, IL-1β is consistently low or undetectable before and after antibiotic administration^6^. This finding argues against a classic “cytokine storm” dominated by IL-1β in human leptospirosis. In contrast, the present study and a previous *in vitro* infection of healthy human whole blood with leptospires both resulted in measurable IL-1β production^24^, likely because whole blood leukocytes retain full inflammasome/pyroptosis responsiveness in the absence of *in vivo* regulatory networks^33^.

Upon leptospirosis, patients exhibit low baseline levels of pro-inflammatory cytokines infection, followed by a sharp inflammatory surge upon β-lactam treatment^6^, highlighting a shared temporal pattern of immune activation between human and mice. This information is important considering the debate on the role of anti-inflammatory drugs in managing spirochaetal JHR^34^. Corticosteroids were among the first agents tested, based on the rationale that dampening the inflammatory cascade could mitigate symptoms. In leptospirosis, the use of steroids is also controversial. Observational studies, including early clinical cohorts of patients with severe pulmonary involvement or Weil’s disease, the most acute form of leptospirosis, have suggested potential benefit from methylprednisome administration, in reducing mortality and improving outcomes in pulmonary complications^35^. However, randomised trial data are limited and inconclusive, and systematic reviews have highlighted methodological constraints and uncertain efficacy^36,37^. Further studies using our models are ongoing to determine whether blocking the inflammation with corticoids has a positive or adverse effect on the severity of infection after antibiotic treatment.

Consistent with the role of β-lactam antibiotics that affects the bacterial cell wall integrity by binding the so-called penicillin binding proteins involved in peptidoglycan synthesis, in 1949, Babudieri *et al.*, already observed by electron microscopy the effect of penicillin on *Leptospira* integrity as they stated that “penicillin dissolves *Leptospira*”^38^. Here we did not visualise the lysis of leptospires upon β-lactam antibiotics but specifically correlated the rapid cytokine production with the efficient killing of leptospires. Further studies are required to investigate the mechanisms and kinetics of leptospiral cell wall degradation by different ß-lactam and other class of antibiotics.

At the mechanistic level, evidence from our study places TLR signalling at the centre of JHR effect initiation. Spirochetes are diderm that share a distinctive cell wall architecture characterised by a periplasmic flagella and high density of lipoproteins, which together shape innate immune recognition and TLR-mediated signalling. In leptospirosis, extensive work has established TLR2 as the dominant receptor mediating host recognition of leptospiral lipoproteins, often tightly associated with LPS^22^ or peptidoglycan^39^. Our data extend these findings by demonstrating that β-lactam-induced JHR effect is critically dependent on TLR2 engagement in both human and mice, consistent with the abrupt exposure of lipoproteins following antibiotic-mediated disruption of the leptospiral envelope. We also identified a role for TLR5 in human whole blood infected with *Leptospira*, implicating flagellin exposure as an additional amplifier of inflammation during JHR. This finding must be interpreted in the context of the ability of pathogenic *Leptospira* to evade TLR5 recognition during infection with live bacteria. Indeed, leptospiral flagella which are confined within the periplasmic space are therefore inaccessible to TLR5, allowing live bacteria to escape flagellin-driven TLR5 innate immune sensing. However, heating or degrading *Leptospira* with antimicrobial peptides, allows for release and exposure of FlaB flagellins, and their recognition by TLR5, although a species specificity has been shown, since human and bovine TLR5 but not murine TLR5 recognised leptospiral flagellins^21^. Accordingly, our data support the concept that β-lactam-mediated disruption of the leptospiral cell wall overcomes this immune evasion strategy, resulting in sudden exposure of normally hidden flagellar components and unveiling a potent TLR5-dependent inflammatory pathway in humans. This mechanism provides a consistent explanation for the cytokine surge observed during the JHR and is fully consistent with the recent clinical study of leptospiral JHR^6^. Another species-specific difference in TLR involvement further refined this model. While human TLR4 is unable to sense leptospiral LPS due to structural differences in lipid A, the mouse TLR4 can signal the presence of leptospiral LPS^9^. Consistently, we did not find a role for TLR4 in JHR-like effect in human blood, although both TLR2 and TLR4 contributed to the inflammatory response during β-lactam treatment in mice, reflecting the unique ability of murine TLR4 to signal through the MyD88 pathway in response to leptospiral LPS^9,22^. The accordance between human *in vitro* and murine *in vivo* data underscores that, despite differences in receptor usage, the downstream consequence of β-lactam treatment is conserved in leptospirosis.

A critical question is whether the JHR severity differs according to leptospiral species or serovars. In leptospirosis, the concept of species- and serovar-specific disease manifestations has long been recognised, with certain strains such as *L. interrogans* serovars Icterohaemorrhagiae, Copenhageni and Manilae, frequently associated with more severe clinical courses than others, reflecting the diverse epidemiology and pathogenic potential of different species described in the literature^40^.

Additional experiments performed with *Leptospira* species representing distinct phylogenetic clades^59^ further refined our understanding of the determinants of the JHR. Strikingly, only low passages of highly pathogenic and virulent *Leptospira interrogans* (P1+) serovars, initially isolated from human patients, consistently induced a robust JHR-like effect following β-lactam treatment. In contrast, saprophytic strains from the S1 (*Leptospira biffexa*) and S2 (*Leptospira kobayashii*) clades, as well as the intermediate P1- (*Leptospira adleri*) and the P2 (*Leptospira licerasiae*) species failed to elicit a detectable reaction. Notably, the lack of an observable JHR effect is not due to an absence of inflammatory response, as these non-P1+ species induced similar levels of pro-inflammatory cytokines in human blood with or without antibiotic treatment. This suggests that they can be sensed prior to antibiotic treatment. However, a broader species/serovar/strain coverage will be important to determine the generalisation of these findings across epidemiological and veterinary contexts. By contrast, highly virulent *Leptospira interrogans* serovars appear to maintain a low inflammatory profile during early infection, likely reflecting their ability to evade or subvert innate immune detection. In this context, antibiotic-mediated bacterial disruption may lead to the sudden exposure of previously concealed immunostimulatory components, triggering a rapid and amplified host response. These findings suggest that the JHR is not a universal feature of leptospiral infections, but rather a phenomenon tightly linked to virulence, host adaptation, and immune evasion strategies. Importantly, the restriction of JHR to clinically derived, highly pathogenic serovars underscores its relevance to human disease and suggests that both bacterial origin and evolutionary niche-whether environmental, reservoir-associated, or human-adapted-critically shape host inflammatory responses. Together, these data further support a model in which JHR emerges from the interaction between antibiotic class, bacterial virulence, and the baseline state of host immune activation, with direct implications for therapeutic strategies and risk stratification in leptospirosis.

In conclusion, this study establishes leptospiral JHR as a treatment-induced, TLR-mediated inflammatory syndrome, where β-lactam antibiotics transform *Leptospira* from stealth pathogens into potent innate immune activators, triggering harmful inflammation. The settings of our mouse and human whole blood models did not allow us to test whether other class of antibiotics would bypass the JHR such as macrolide, like azithromycin or doxycycline, a tetracycline, already preferred in endemic countries like Thailand for its accessibility and tolerability^32^. Further studies would be required to potentially underscoring its potential as a safer alternative.

Preventing JHR must be a clinical priority, not an afterthought, given its predictable and mechanistically driven nature. Future clinical studies should investigate the risks/benefits and choice of treatments in patients in different contexts of health care systems and circulating *Leptospira* strains.

## Methods

### Leptospira interrogans culture

The pathogenic *Leptospira interrogans* (*L. i*) serovars used in this study were the Manilae strain MFLum1^20^ (a bioluminescent derivative of strain L495), the Icterohaemorrhagiae strain Verdun (virulent Cl3)^8,41^, and the Copenhageni strains LV2756^41^. These strains were originally isolated from human patients with leptospirosis in the Philippines, France, and Brazil, respectively. Non-virulent derivatives included the *L. i* Icterohaemorrhagiae Verdun Avir p104 strain^41^ and the *L. i* Copenhageni Fiocruz L1-130 mutant FGLum4 strain^42^. Other strains used included *L. adleri* FH2-B-D1 and *L. licerasiae* strain VAR 010 *strains*, known to be less virulent or intermediate *Leptospira* species^43^. We also used the non-pathogenic saprophytic *Leptospira biffexa* serovar Patoc strain Patoc I and its bioluminescent derivative PFLum7^20^ and *L. kobayashii* E30 strains.

All strains were cultured in liquid Ellinghausen-McCullough-Johnson-Harris (EMJH) medium at 30 °C without agitation (Table 1). The bioluminescent strains MFLum1, PFLum7 and FGLum4 were previously generated by random transposon mutagenesis introducing a luciferase cassette^20^. MFLum1 retained the growth characteristics and overall pathogenicity of the parental strain. *Leptospira* cultures were passaged weekly (twice weekly for *L. biffexa*) and adjusted to a final concentration of 10^6^ bacteria/mL. Bacteria were used one week later, at the end of the exponential growth phase. For infection experiments, leptospires were centrifuged at 3 200 x g for 25 min at room temperature, resuspended in endotoxin-free phosphate-buffered saline (PBS, Gibco), and enumerated using a Petroff-Hausser counting chamber. *L. interrogans* Manilae MFLum1, Icterohaemorrhagiae Verdun Cl3, Copenhageni Fiocruz LV2756, *L. adleri, L. licerasiae* and *L. kobayashii* were used between passages 3 and 12. The laboratory *L. biffexa* Patoc was high passaged strain.

### Borrelia burgdorferi culture

The pathogenic *Borrelia burgdorferi* sensu stricto strain B31-A3 was cultured in liquid Barbour-Stoenner-Kelly (BSK) medium supplemented with rabbit serum at 30 °C without agitation (Table 1). For each experiment, a fresh frozen aliquot was thawed. For cellular infection, bacteria were centrifuged at 3 200 x g for 25 min at room temperature, resuspended in endotoxin-free PBS (PBS, Gibco), and counted using a Petroff-Hausser chamber.

### Blood donors

Human peripheral blood samples were collected from healthy volunteers through the Clinical Investigation Centre INVOLvE (Investigation and volunteers for human health) at Institut Pasteur. The participants received oral and written information about the research (authorised project LEPTBLOOD 2023-029) and gave written informed consent in the frame of the COSIPOP cohort after approval of the CPP Est II Ethics Committee (2023, February 20^th^).

### Whole-blood infection assays

Whole-blood samples were collected from healthy male and female volunteers into heparinised tubes and processed immediately. Aliquots of whole blood were dispensed into 96-well plates and infected with 10^7^ or 10^6^ live *Leptospira* or *Borrelia* in a final volume of 190 µL. PBS was used as a negative control. At 5 h post-infection was added, 10 µL of amoxicillin at final concentrations of 5 µg/mL, or ceftriaxone, at 1 µg/mL. Cultures were incubated at 37 °C for 24 h before assessment of bacterial viability and cytokine production.

### Neutralisation of Toll-like receptors in whole blood

For Toll-like receptor (TLR) neutralisation, whole blood (150 µL) was incubated for 1 h at 37 °C with 50 µL of neutralising antibodies against TLR2, TLR4, or TLR5 (InvivoGen) at a final concentration of 10 µg/mL, or with the corresponding isotype controls (IgA2 or IgG). Blood was then infected with 10^6^ or 10^5^ *L. interrogans*. After 5 h, 10 µL of amoxicillin was added at 5 µg/mL. Lipopolysaccharide (LPS, 10 ng/mL), Pam3CSK4 (100 ng/mL), and flagellin (50 ng/mL) were used as positive controls for TLR4, TLR2, and TLR5 neutralisation, respectively. Bacterial viability and cytokine concentrations were assessed 24 h post-infection (Table 2).

**Table 2.** Neutralisation antibodies of TLRs and agonists (InvivoGen).

| Receptor | Agonist + Cat. No. | Neutralising Antibody + Cat. No. | Clone | Isotype control + Cat. No. |
| --- | --- | --- | --- | --- |
| TLR2 | Pam3CSK, *tlrl-pms* | Anti-hTLR2-IgA2, *htlr2-mab7-2* | B4H2 | Anti-β-Gal-hIgA2, *bgal-mab7* |
| TLR4 | Ultrapure LPS O111:B4, *tlrl-3pelps* | Anti-hTLR4-IgG, *mabg-htlr4-2* | W7C11 | Anti-β-Gal-mIgG1, *bgal-mab9* |
| TLR5 | Ultrapure flagellin, *tlrl-epstfla-5* | Anti-hTLR5-IgA2, *htlr5-mab7-2* | Q2G4 | Anti-β-Gal-hIgA2, *bgal-mab7* |

### *In vivo* infection and antibiotic treatment in mice

Adult (6-10-week-old) wild-type C57BL/6J mice were obtained from Janvier Labs and acclimatised for at least one week at the Institut Pasteur animal facility. On arrival at the animal facility of Institut Pasteur, the mice were randomly placed in cages with nesting aids and allowed to acclimatise for 1 week. *Tlr2^-/-^* and *Tlr4^-/^* mice in a C57BL/6J background were bred and maintained in-house. Groups of three to five mice of each sex were intraperitoneally infected with 10^8^ *L. interrogans* serovar Manilae MFLum1 in 200 µL PBS. Control mice received PBS alone. Body weight, temperature, and clinical signs were monitored daily. Clinical scores were calculated as a composite of weight loss, alertness, locomotion, and body condition, as described previously^5^. On day 3 post-infection (pi), corresponding to the peak of leptospiral dissemination, mice received intraperitoneal injections of 100 µL of amoxicillin (100 µg/mouse)^20^. Mice were euthanised 3 h after antibiotic administration by cervical dislocation or progressive CO_2_ inhalation. Blood was collected by cardiac puncture into serum-separation tubes (Microvette 200 serum-gel, Sarstedt), centrifuged, and serum was aliquoted and stored at −20 °C. A total of 110 mice, 60 males, and 50 females, were used in this study.

### Immunohistochemistry

*Staining:* Hearts were harvested on day 3 pi and fixed directly in 10% neutral buffered formalin for 48 h. Fixed tissues were further washed in 70% ethanol and embedded in paraffin. Immunohistochemical studies were performed using antibodies against macrophages (F4/80) and T cells (CD3) (Table 3). Stained slides were evaluated using a ZEISS AxioScan.Z1 digital slide scanner and Zen software (Zeiss).

**Table 3.** Antibodies used for immunohistochemistry.

| Anti-mouse | Target cells | Dilution | Reference | RRID |
| --- | --- | --- | --- | --- |
| Anti-F4/80 rabbit mAb | Macrophages | 1/300 | D2S9R, Cell signalling | AB_2799771 |
| Anti-CD3 rabbit pAb | T cells | 1/400 | A0452, Dako | AB_2341188 |

*Image analysis:* Quantitative image analysis was performed using Visiopharm (version 2025.02 x64). Tissue regions were automatically identified using the Tissue Detection application to exclude background and artefactual areas. Nuclear segmentation was performed using the Nuclei Detection, AI (Brightfield) algorithm. Detection accuracy was optimised by applying thresholds to the “Nuclei” and “HDAB - Haematoxylin” features derived from H-DAB colour deconvolution, allowing removal of non-specific objects. Nuclei were then classified as positive or negative based on DAB staining intensity. Intensity variation thresholds were applied to distinguish marker-positive (CD3 or F4/80) from marker-negative cells.

*Quantification:* The following parameters were extracted for each sample: total nuclei, positive nuclei, negative nuclei, positive fraction, and percentage of positive nuclei. All analyses were performed using identical parameter settings within each experiment.

### Bacterial survival assays

Bacterial survival was assessed by inoculating 100 µL of whole blood or culture supernatants into 2 mL EMJH medium for non-bioluminescent strains (Verdun and Fiocruz LV2657) and incubating cultures at 30 °C for 7 days before measuring optical density at 420 nm. For the bioluminescent strain MFLum1, samples were transferred directly into white 96-well plates and luminescence was measured following addition of D-luciferin (667 µg/mL, XenoLight D-Luciferin-K⁺ Salt, PerkinElmer) prepared in endotoxin-free PBS (Gibco). Luminescence was quantified using a luminometer, and absorbance was measured using a TECAN microplate reader.

### Cytokine and biochemical marker measurements

Whole blood and mouse serum collected on day 3 pi were stored at −20 °C until analysis. Concentrations of TNF, IL-1β, IL-6, IL-10, and CCL5/RANTES were quantified using ELISA kits (RCD Systems) according to the manufacturer’s instructions. Serum amyloid A (SAA) and cardiac troponin I (cTnI) concentrations were measured using ELISA kits (Abcam) (Table 4).

**Table 4.**
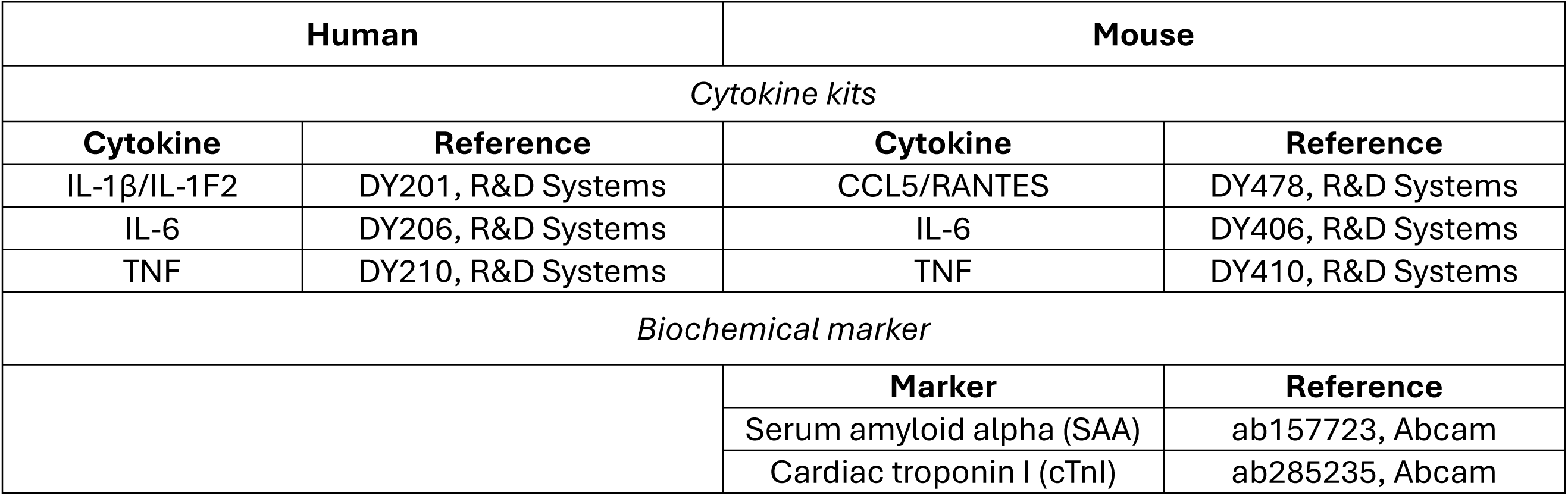
Kits used for cytokine and biochemical parameters quantification.

### Lactate dehydrogenase assay

Lactate dehydrogenase (LDH) release was quantified in serum and homogenised organs (liver, kidney, lung, and spleen) collected on day 3 pi. Samples were diluted to 500 µg/mL and analysed using the CyQUANT LDH Fluorometric Assay (Invitrogen) following the manufacturer’s protocol. Fluorescence was measured at 560 nm excitation and 590 nm emission using a TECAN Spark fluorimeter.

### Assessment of vascular permeability

On day 3 pi, 2,5 hours after antibiotic administration. Anaesthesia was induced and maintained with 2.5% isoflurane using an XGI-8 induction chamber for 10 mins, and anaesthetised mice received an intravenous retro-orbital injection of 100 µL sterile 0.5% (w/v) Evans blue dye (Sigma) in PBS^5^. After 30 min, mice were euthanised with progressive CO2 inhalation and perfused with 20 mL PBS containing heparin (2 U/mL). Organs were collected, weighed, and incubated in 500 µL formamide (Sigma) at 37 °C for at least 48 h. After centrifugation, absorbance of the supernatant was measured at 620 nm, and Evans blue concentrations were calculated.

### Reagent validation

All antibodies used in the present study were commercially available. The RRID tags of the antibodies used for the immunohistochemical staining are given in Table 3. For the neutralisation of the TLRs, the RRID tags were not available for the antibodies used, but instead the exact clone and the reference number of the products are indicated in Table 2.

### Ethics

All animal experiments complied with ARRIVE guidelines and European and French regulations for animal research and were approved by the Institut Pasteur Animal Welfare Committee (CETEA, protocol dap210024 / APAFIS#30877-2021040212335074 v1).

### Statistics

Statistical analyses were performed using GraphPad Prism (version 10.6.1). For *in vitro* experiments using human whole blood, cytokine responses were analysed using paired two-way ANOVA with Šidák’s multiple comparison test. The two factors included (1) technical replicates within each donor and (2) treatment condition (e.g., antibiotic or neutralising antibody). Technical replicates were included in the statistical model to account for within-donor variability. However, graphical representations display one data point per donor corresponding to the mean of technical replicates, to reflect biological variability between individuals. For *in vivo* mouse experiments comparing two groups, Welch’s t test was used. For time-course experiments or analyses involving multiple factors (e.g., genotype and treatment), two-way ANOVA with Šidák’s multiple comparison test was applied as specified in figure legends. Data are presented as mean values unless otherwise indicated. A two-sided *p* value ff 0.05 was considered statistically significant.

### Contributors

SP: investigation, data analysis and interpretation, figures, supervision of students, methodology, writing original draft-review editing, TB: investigation, DJ: investigation, JC: preliminary experiments, FVP: investigation, resources (*Leptospira* cultures and mice ethical authorisation), review editing, HL: resources (supervision healthy volunteers cohort, and blood sampling), CW: conception, grant acquisition, investigation, data analysis and interpretation, supervision, visualisation, writing draft-review editing

### Data sharing statement

All data are available in the main text or in the Supplementary Material. Raw data and analyses are secured on elabnotebook at Institut Pasteur and will be made available on request to Catherine Werts.

## Acknowledgments

We are grateful to the healthy volunteers of the COSIPOP cohort who agreed to the scientific use of their samples and data. We thank the CRBIP unit CHIP for providing fit-for-purpose biological resources and associated services. We thank Dr. Mathieu Picardeau (Unité Biologie des Spirochètes, Institut Pasteur) and Dr Roman Thibeaux (Leptospirosis Research and Expertise Unit, Institut Pasteur of New Caledonia) for kindly providing us with the bacteria. We thank Elise Jacquemet (Hub of bioinformatic and biostatistics, Institut Pasteur) for help in statistics analysis. We thank members of the Histopathology Core Facility at Institut Pasteur, and specifically, David Hing and David Hardy, for sample preparation, staining and analysis of mice tissues, and Sarra Loulizi for quantification of heart tissues. We thank Dr. Patrick Lefèvre, Dr. Cécile Cazorla (CHT, Medipole, New Caledonia), and Dr. Anne-Françoise Loarec (Medical epidemiology unit, Institut Pasteur of New Caledonia), for helpful discussions about leptospirosis, treatments and JHR. We thank Dr. David Haake (UCLA, USA) for his kind support and for including our group in the NIH project. We thank Pr. Jean-Marc Cavaillon for critical reading of the manuscript. We thank Dr. Ivo Gomperts Boneca (BGPB Unit, Institut Pasteur) for constant support and hosting our group. We used DeepL write software for English editing.

## Supplementary Material

**Supplementary Figure 1.**
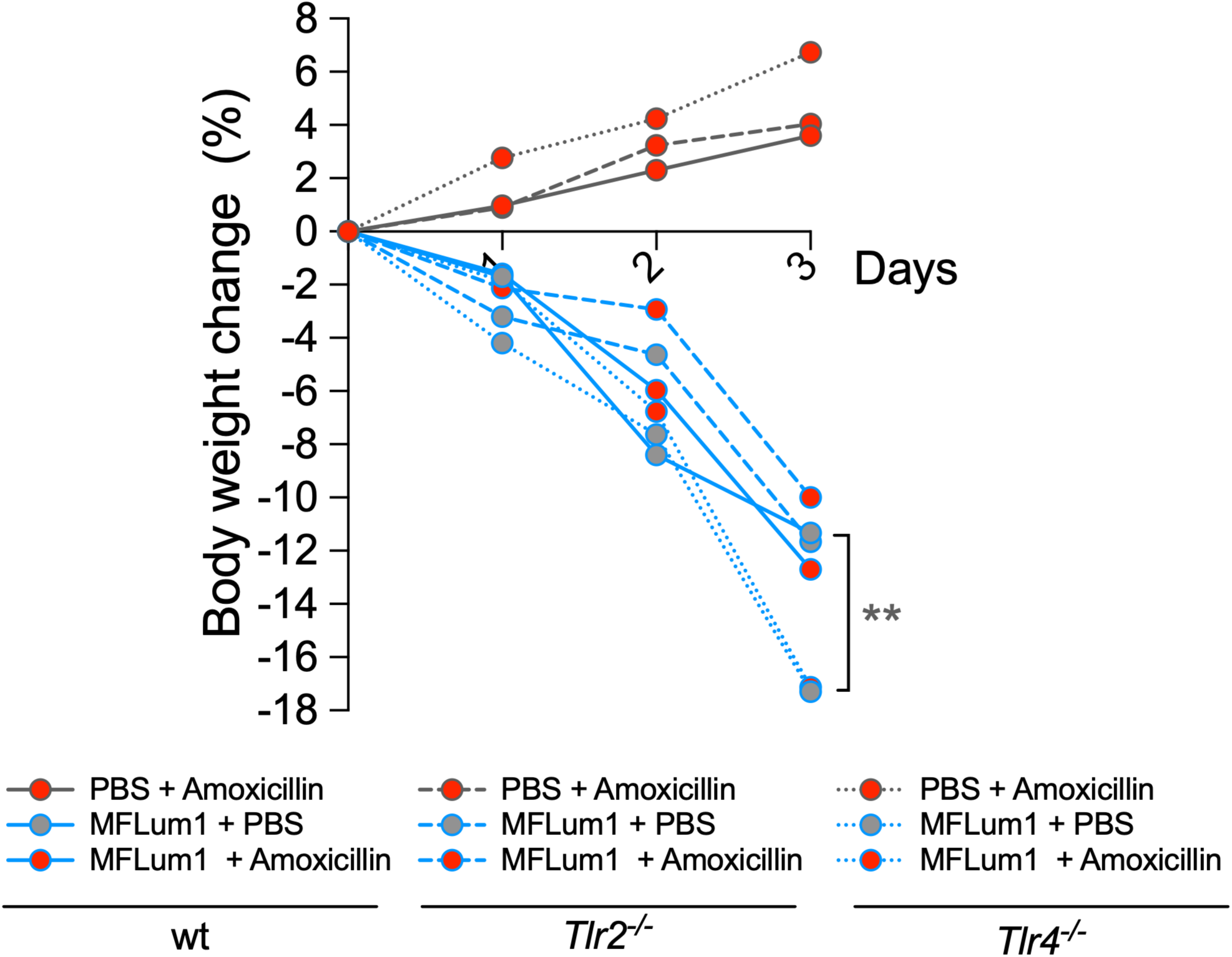
Weight loss of infected WT and TLR-deficient mice receiving amoxicillin. Body-weight changes in non-infected or infected wild-type, *tlr2*-/-, and *tlr4*-/- mice following amoxicillin administration. Continuous line indicates wild-type mice; dashed line indicates *tlr2*-/- mice; dotted line indicates *tlr4*-/- mice. Grey line with red symbols indicates uninfected mice treated with amoxicillin; blue line with grey symbols indicates infected untreated mice; blue line with red symbols indicates infected mice treated with amoxicillin. Data are from one representative experiment of three performed. Statistical analyses were performed using two-way ANOVA with Šidák’s multiple comparison test (factor 1: day post-infection; factor 2: genotype). Only significant comparisons between infected treated groups are shown. Significance threshold was p ff 0.05 (*p ff 0.05, **p ff 0.01).

**Supplementary Figure 2.**
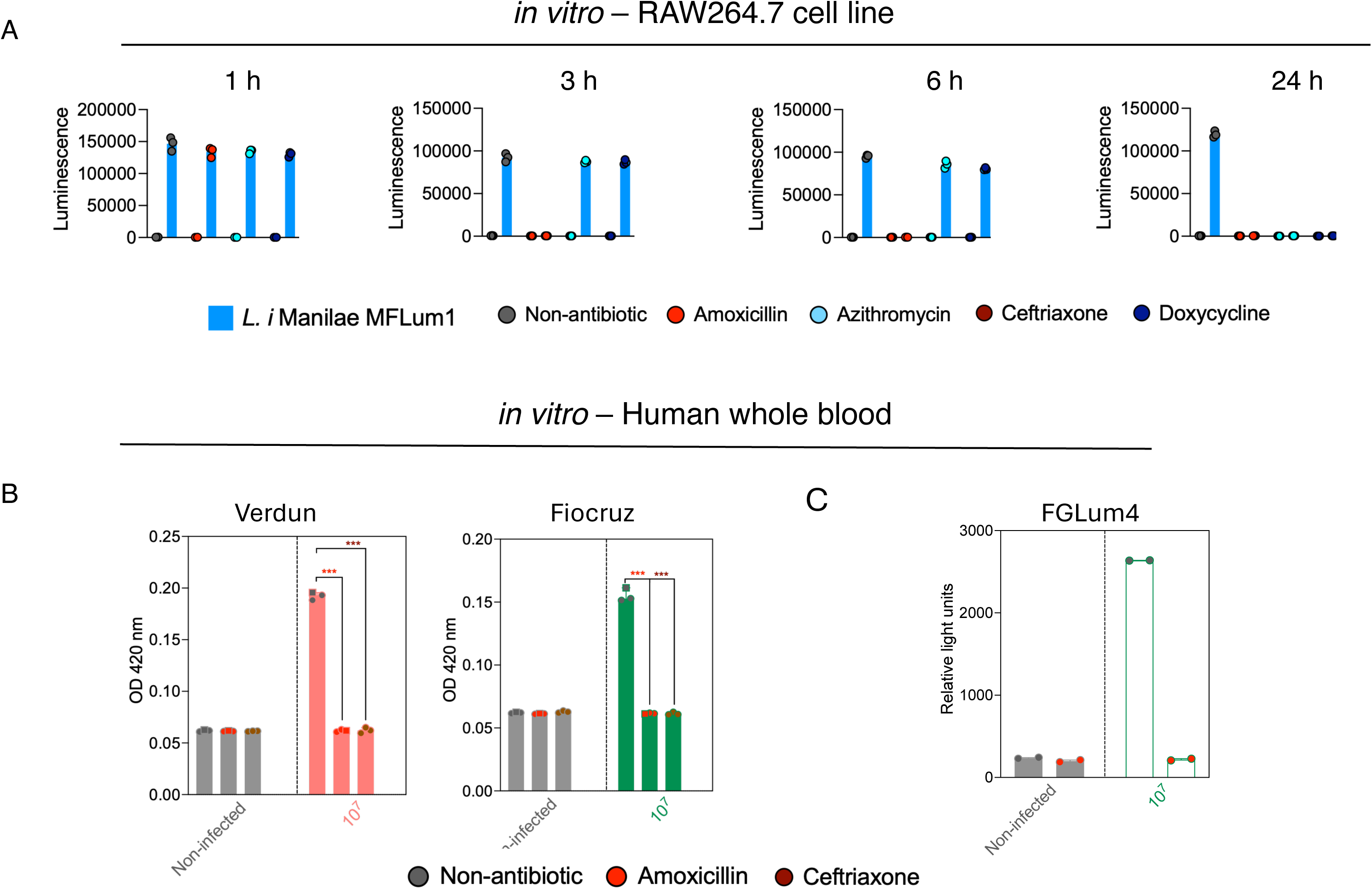
**Antibiotic killing efficiency *in vitro*** A) direct measures by bioluminescence of bacterial killing by antibiotics. Kinetics of antibiotic effects on leptospiral survival measured 1, 3, 6 and 24 hours post antibiotic treatment by bioluminescence in murine macrophage cell line RAW.264.7 infected with *Leptospira interrogans* serovar Manilae bioluminescent strain MFLum1 for 3h before antibiotic treatment. Grey symbols indicate untreated infected cells; red symbols indicate amoxicillin; light blue symbols indicate azithromycin; brown symbols indicate ceftriaxone, and dark blue symbols indicate doxycycline. B) Indirect measures of bacterial killing. Optical density at 420 nm of *Leptospira interrogans* serovar Icterohaemorrhagiae and Copenhageni inoculated in EMJH for 7 days 24 h post infection in whole human blood and treated with amoxicillin, or ceftriaxone. Grey symbols indicate infected untreated blood; red symbols indicate amoxicillin; brown symbols indicate ceftriaxone. Data are mean of n=3 donors pooled from three independent experiments. Statistical analyses were performed using two-way ANOVA with Šidak’s multiple comparison test (factor 1: technical replicates per donor; factor 2: antibiotic regimen). C) Direct measures by bioluminescence of *L. interrrogans* non-virulent strain Copenhageni Fiocruz FGLum4 without (grey symbols) of after treatment with Amoxicillin (red symbols).

**Supplementary Figure 3.**
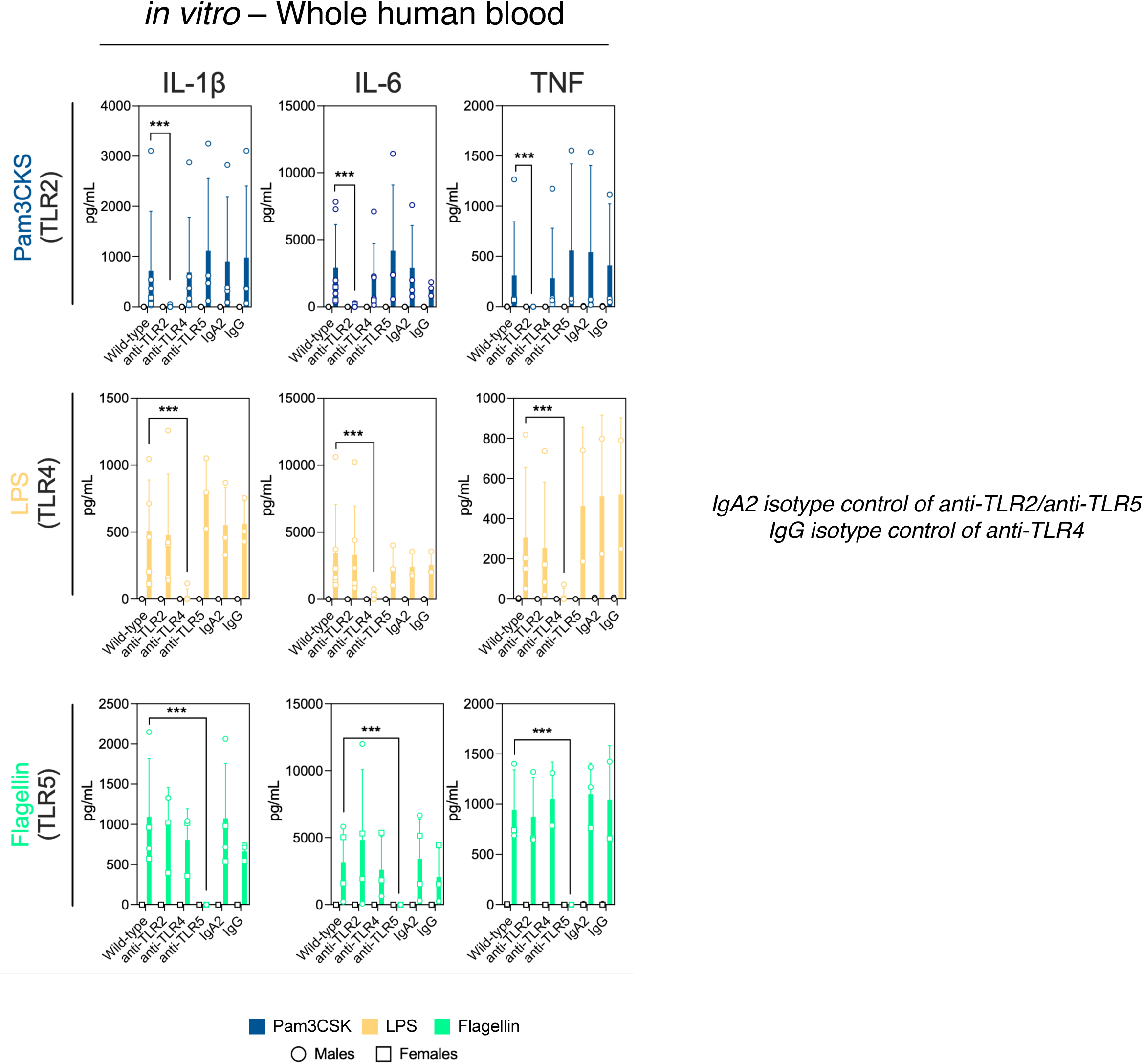
Validation of TLR neutralisation efficiency in whole blood. In vitro stimulation of human whole blood with TLR agonists to verify the respective receptor neutralisation. Pam3CSK4 (blue) was used as a TLR2 agonist; lipopolysaccharide (LPS; yellow) as a TLR4 agonist; and flagellin (bright green) as a TLR5 agonist. Cytokine concentrations (IL-6, TNF, and IL-1β) were measured in whole blood stimulated with each agonist in the absence of neutralising antibodies (wild-type condition) or in the presence of neutralising antibodies against TLR2, TLR4, or TLR5, or corresponding isotype controls (IgA2 or IgG). Data are mean of n=2-6 donors pooled from two to six independent experiments. Statistical analyses were performed using two-way ANOVA with Šidák’s multiple comparison test (factor 1: technical replicates per donor; factor 2: neutralising antibody or isotype condition). Only significant comparisons between stimulated untreated and antibody-treated conditions are shown. Open circles represent male donors, and open squares represent female donors. Significance threshold was p ff 0.05 (*p ff 0.05, **p ff 0.01, ***p ff 0.001).

**Supplementary Figure 4.**
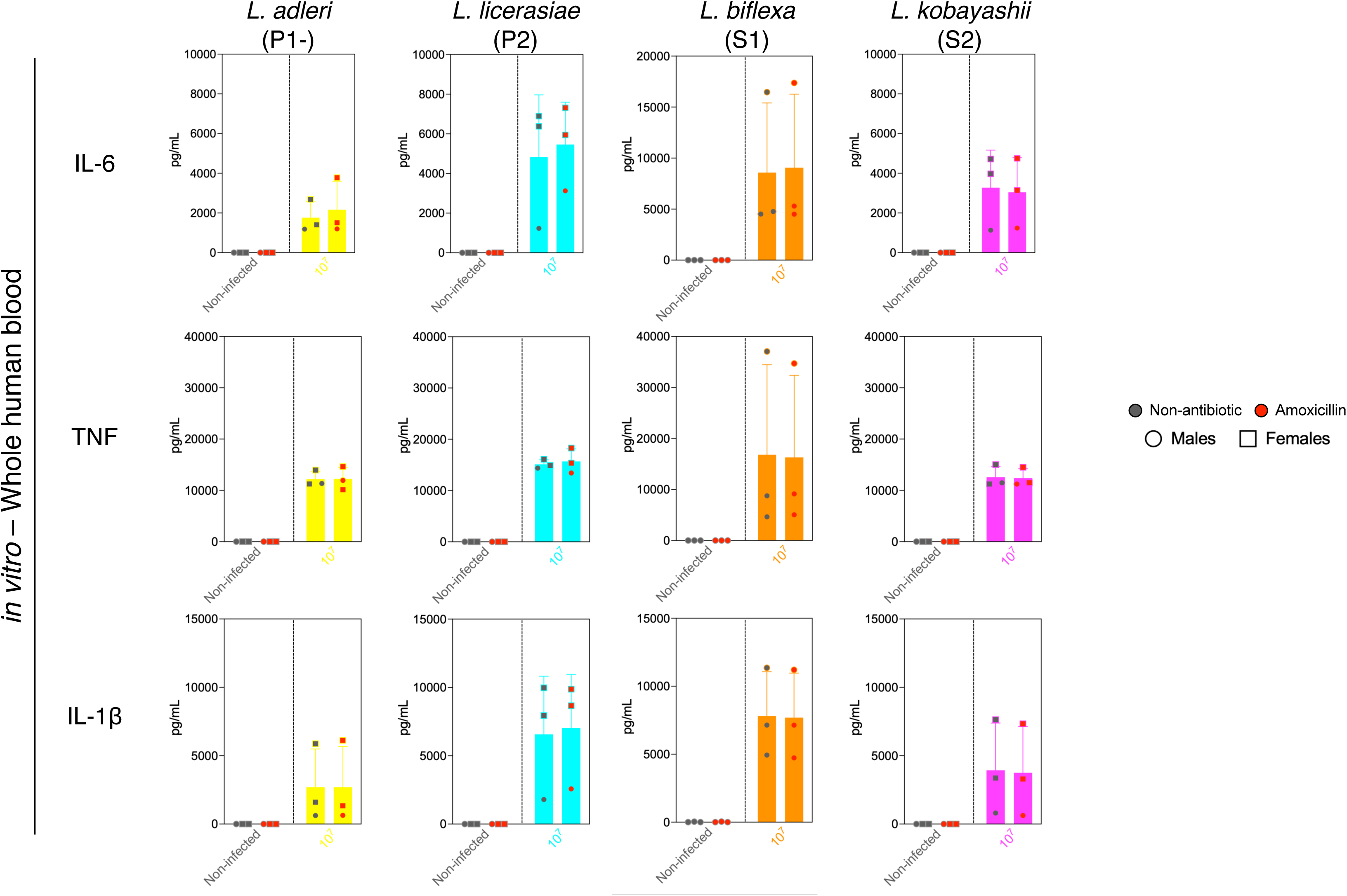
***Leptospira* species other than *Leptospira interrogans* do not induce a Jarisch-Herxheimer reaction.** Comparison of *in vitro* inflammatory responses in human whole blood infected with different *Leptospira* species following amoxicillin treatment. From left to right: *Leptospira adleri (*P1-), *Leptospira licerasiae* (P2), *Leptospira biffexa* (S1), and *Leptospira kobayashii* (S2). Cytokine concentrations (IL-6, TNF, and IL-1β) were measured in infected whole blood following treatment with amoxicillin (red symbols) or in untreated conditions (grey symbols). Data are shown for n=3 independent experiments.

